# Cost utility analysis of end stage renal disease treatment in Ministry of Health dialysis centres, Malaysia: hemodialysis versus continuous ambulatory peritoneal dialysis

**DOI:** 10.1101/660167

**Authors:** Naren Kumar Surendra, Mohd Rizal Abdul Manaf, Lai Seong Hooi, Sunita Bavanandan, Fariz Safhan Mohamad Nor, Shahnaz Shah Firdaus Khan, Ong Loke Meng, Abdul Halim Abdul Gafor

## Abstract

**OBJECTIVES:** In Malaysia, there is exponential growth of patients on dialysis. Dialysis treatment consumes a considerable portion of healthcare expenditure. Comparative assessment of their cost effectiveness can assist in providing a rational basis for preference of dialysis modalities.

**METHODS:** A cost utility study of hemodialysis (HD) and continuous ambulatory peritoneal dialysis (CAPD) was conducted from a Ministry of Health (MOH) perspective. A Markov model was also developed to investigate the cost effectiveness of increasing uptake of CAPD to 55% and 60 % versus current practice of 40% CAPD in a five-year temporal horizon. A scenario with 30% CAPD was also measured. The costs and utilities were sourced from published data which were collected as part of this study. The transitional probabilities and survival estimates were obtained from the Malaysia Dialysis and Transplant Registry (MDTR). The outcome measures were cost per life year (LY), cost per quality adjusted LY (QALY) and incremental cost effectiveness ratio (ICER) for the Markov model. Sensitivity analyses were performed.

**RESULTS:** LYs saved for HD was 4.15 years and 3.70 years for CAPD. QALYs saved for HD was 3.544 years and 3.348 for CAPD. Cost per LY saved was RM39,791 for HD and RM37,576 for CAPD. The cost per QALY gained was RM46,595 for HD and RM41,527 for CAPD. The Markov model showed commencement of CAPD in 50% of ESRD patients as initial dialysis modality was very cost-effective versus current practice of 40% within MOH. Reduction in CAPD use was associated with higher costs and a small devaluation in QALYs.

**CONCLUSIONS:** These findings suggest provision of both modalities is fiscally feasible; increasing CAPD as initial dialysis modality would be more cost-effective.

## 1.0 Introduction

Renal replacement therapy (RRT) is the usual choice of treatment for patients suffering from end stage renal disease (ESRD), which includes dialysis, either hemodialysis (HD) or peritoneal dialysis (PD) and a kidney transplant. A kidney transplant is the best choice of treatment in patients suffering from ESRD, however, the waiting list for transplantation continue to grow despite kidney transplants from live donors due to the organ scarcity [1].

Dialysis modality selection in various countries is influenced by non-medical factors including financial and reimbursement policy [2-4]. Although both HD and PD are costly, specific advantages and disadvantages have been identified for each of them. Comparative assessment of their cost effectiveness can assist in providing a rational basis for preference of one or the others [5]. Economic evaluation of ESRD treatment and policy explorations have been performed recurrently in many settings [6]. However, economic evaluations of dialysis modalities in Malaysia are still lacking despite the continuous growth of ESRD patients at an alarming rate. Peritoneal dialysis is underutilized although it is considered a more cost-effective, if not, equally cost-effective treatment as compared to HD around the world [1, 7-9].

Dialysis provision is dominated by HD in Malaysia and there is an inequitable distribution of its provision. Dialysis acceptance rates have reached a level equal to that of developed countries [1, 10]. According to the 24^th^ report of the Malaysian Dialysis and Transplant Registry (MDTR), 6,662 new HD patients and 1,001 new PD patients were reported in 2016 representing an acceptance rate of 216 per million population (pmp) and 32 pmp respectively. Overall, the total number of HD and PD patients increased to 35,781 patients (1,159 pmp) and 3,930 patients (127 pmp) respectively in 2016 [11]. The number of dialysis centres for the whole of Malaysia increased from 698 in 2011 to 814 in 2016. This was attributed by the private dialysis centres which had trebled from 5 pmp in 2004 to 14 pmp in 2016 [11].

ESRD has significant economic consequences with loss of gross domestic product (GDP) for its management. In developed countries, it was reported that the expenses for RRT provision were 2-3% of total healthcare expenditure while ESRD patients accounted for just 0.02-0.03% of the total population [12]. Although limited data is available for ESRD expenditure in Malaysia, the estimated costs of dialysis in 2005 were RM379.1 mil [1, 10]. A recent forecast estimates the cost incurred to treat 51,269 patients with dialysis in the year 2020 is RM1.5 billion (USD384.5 million) [13]. Given the low organ donation rate and continual growth of ESRD population, it is timely to carry out an economic evaluation of HD and PD.

The aim of this study is to compare the cost utility of HD and CAPD and to assess the cost utility of different dialysis provision strategies at varying levels of CAPD usage versus current practice using a Markov model simulation cohort.

## 2.0 Methods

The study used both primary and secondary data for HD and CAPD. The primary outcomes of interest were costs and utilities of HD and CAPD derived from the primary data collection as part of this study and these have been published [15, 16]. The survival data was sourced from the Malaysian Dialysis and Transplant Registry (MDTR). The perspective of the study was that of the MOH because it is the ultimate decision maker on the funding of its own dialysis programme. Sources of data used in the study are as shown in Table 1.

**Table 1:** Sources of data

| Data | Data Type | Source |
| --- | --- | --- |
| Cost | Primary data | Surendra et al. 2018 [14] |
| Utilities (EQ-5D) | Primary data | Surendra et al. 2019 [15] |
| Life years (LY) | Secondary data | MDTR |
| Transitional probabilities | Secondary data | MDTR |

\*MDTR-Malaysia Dialysis and Transplant Registry

A Markov model cohort simulation was developed to explore the cost utility of hypothetical dialysis provision strategies versus current practice.

### 2.1 Costs

The mean costs per patient per year were obtained in the cost analysis and the results have been published [14]. The costs were divided into components which include access surgeries, outpatient clinic care, dialysis consumables, staff emoluments, land, building and hospitalizations. All costs were presented in Malaysian Ringgit (RM) valued in the year 2017.

### 2.2 Health utilities

Patient responses to the EQ-5D-3L were used to generate a health state profile that was converted to index-based values. The Malaysian value-set was used, and the results have been published [15].

### 2.3 Survival estimates

The Kaplan-Meier product-limit survivor function approach was used to estimate the mean survival rates (life years) for HD and CAPD patients because it best fits the available data. Transitional probabilities to death and change between the modalities were also estimated. The survival dataset was obtained from the MDTR. The samples were all HD and all CAPD patients who began dialysis in MOH centres between 2011 and 2015. The outcomes of interest are death and change of modality and the follow-up period ended on 31^st^ December 2016.

#### 2.3.1 Life years

Survival was not censored for change of modality based on first modality. Survival durations for patients were calculated from the date commencing the first modality till 31^st^ December 2016 for patients who were still on dialysis. For patients who died, survival duration was calculated from date commencing the first modality, till date of death. All death outcomes whether occurring during first modality or after change in modality were considered for this analysis. Patients were censored if they had received a kidney transplant, recovered kidney function and were lost to follow up during the period.

#### 2.3.2 Transition probability-change of modality

Annual change of modality rates was calculated by dividing the number of the events in a year by the estimated mid-year patient population. The proportion of cohort in each dialysis modality and transitioning between the modalities were imputed based on the observed mean dialysis change rates among HD and CAPD patients over the five years period. The rates were converted into an annual transition probability by using the following formula: p = 1 – exp (-r*t) where p is the per cycle probability, r is the per-cycle rate, and t is the number of cycles. The probabilities were converted using the method on probabilities and rates by Drummond et.al. (2015) [16].

#### 2.3.3 Transition probability-death

Annual death rates were calculated by dividing the number of deaths in a year by the estimated mid-year patient population. The annual transition probabilities from HD to death and from CAPD to death were determined based on the observed mean death rates over the five years period. The rates were converted into an annual transition probability by using the following formula; p = 1 – exp (-r*t) where p is the per cycle probability, r is the per-cycle rate, and t is the number of cycles.

### 2.4 Markov model simulation cohort

The model was developed based on the Markov model designed by Villa et al. (2011) [17]. Only three health states were included in this model; HD, CAPD and death as shown in Figure 1. The theoretical model structure was built in the TreeAge Pro software version 2018 to run a computer-generated simulation on a hypothetical cohort of dialysis patients stating either HD or CAPD. In this study, the model simulated progression of renal outcomes in temporal horizons of five years. Each cycle consumes one year. Thus, this model runs in five cycles.

**Figure 1:**
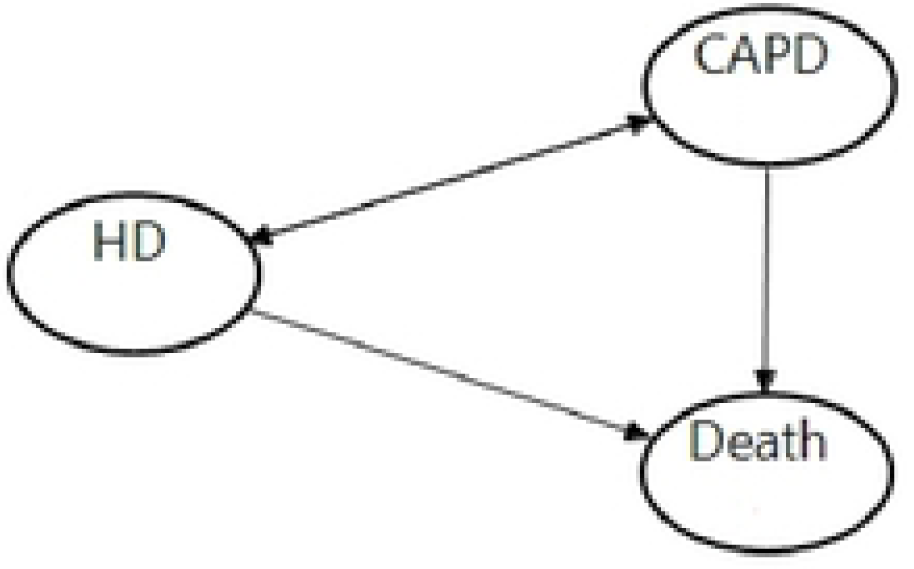
Markov model transition diagram.

#### 2.4.1 Scenario consideration

According to the MDTR data, 60% of all patients dialysing at MOH centres were on HD and 40% were on CAPD. Hence, this observed distribution was used as the base case scenario in this study. Alternative scenarios to Malaysia current practice included: Scenario 1, a model with an increased initial distribution of CAPD by 5%; Scenario 2: a model with an increased initial distribution of CAPD by 10%; Scenario 3: a model with a decreased initial distribution of CAPD by 10%.

#### 2.4.2 Model assumptions

The underlying assumption of a Markov model in its standardized version is independent from past events, the Markovian property [16]. This means that irrespective of which state an individual in the model comes from, the patient will still face the same transition probabilities as someone who has another past state. A half-cycle correction was employed, which is equivalent to an assumption that, state transitions occur, on average, halfway through each cycle. Additionally, the model undertook the following assumptions; a) the Markov cohort comprised of ESRD patients aged 18 years and older, various racial/ethnic groups and clinical characteristics reflecting the characteristics of real world dialysis patients in Malaysia; b) the cohort starts with an initial distribution observed in each scenario; c) ESRD patients with no contraindications to any modality; d) patients’ characteristics (other than age) remain unchanged during each cycle.

#### 2.4.3 Model inputs

Relevant model data were incorporated based on primary data which were collected as part of this study and the detailed methodology and results have been published elsewhere [14, 15].

Transition probabilities were estimated according to an analysis of a de-identified dataset from MDTR as described above. The transition probabilities were assigned to each modality including death. Three health states (HD, CAPD, Death) were defined, with the chance of bidirectional transitions between all the states except death, which is an absorbent state. The total of probability must add up to one in each scenario. The initial prevalence was distributed among the modalities according to the proportions observed in the latest MDTR data. Based on those data, the future prevalence in each cycle (5 year) and state were determined by the application of a transition probabilities matrix (TPM). In the model, from one cycle to the next, the patient may stay on their current modality, switch to a different modality or die. Patients may die in any state (HD or CAPD) and only one movement was allowed per cycle. Once a patient dies, he/she no longer accrue costs and benefits. Table 1 shows the model inputs.

#### 2.4.4 One-way sensitivity analysis

One-way sensitivity analysis was used to investigate variability on all parameters included in the model. The plausible ranges of transition probabilities, health utilities and maximum/minimum value of cost components were included in this analysis. The results were presented in Tornado diagrams based on Net Monetary Benefit (NHB). A Tornado diagram is a special bar chart which is the graphical output of a comparative sensitivity analysis. It is comparing the relative importance of variables considered in the model [16]. The NHB was preferred due to the minute effectiveness differences between the strategies. It is calculated as (incremental benefit x threshold – incremental cost). A positive NHB indicates that the imputed values are cost-effective at the given cost effectiveness threshold.

#### 2.4.5 Probabilistic sensitivity analysis

To evaluate the impact of uncertainty on all the parameter values simultaneously, a probabilistic sensitivity analysis was performed by second order Monte Carlo simulations (1000 iterations). Each simulation provided one value of cost effectiveness. A gamma distribution for costs and a beta distribution for utilities and transition probabilities were used. Costs and outcomes were undiscounted or discounted at an annual rate of 3%. The result is presented in a cost effectiveness acceptance curve (CEAC).

### 2.5 Cost effectiveness threshold

Costs per QALY and LY less than three times and one-time gross domestic product per capita (GDP) are cost-effective and very cost-effective, respectively [18]. In Malaysia, the GDP per capita in 2017 was US$9,660 (≈RM40,000) [19]. Therefore costs per LY or QALY should be lower than RM120,000 per patient to be cost-effective. The combined data of costs and utilities are shown in Table 2.

**Table 2:**
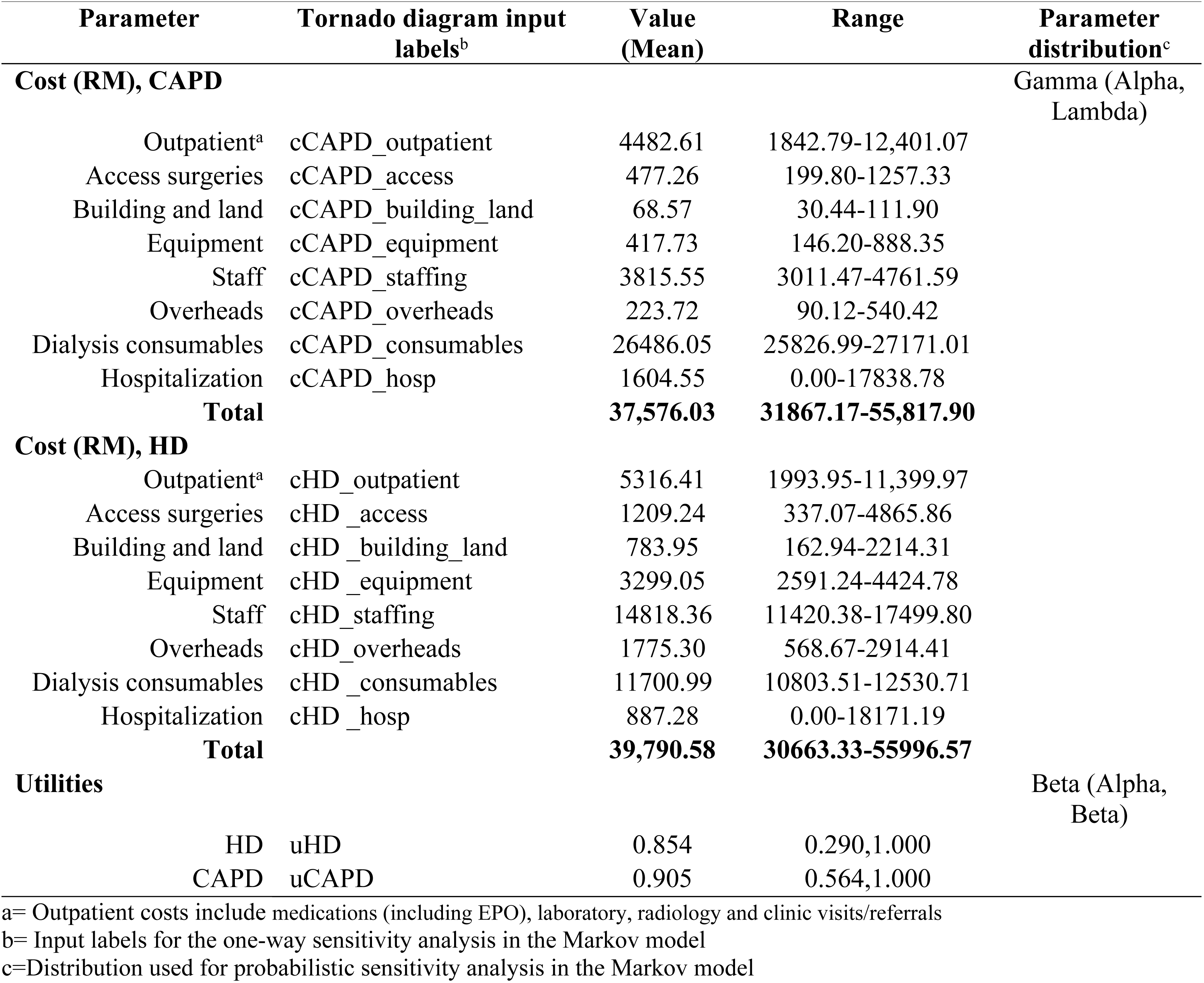
Parameter inputs for Markov model cohort simulation

### 2.6 Incremental cost effectiveness ratio (ICER)

For the Markov model, the primary outcome is the Incremental Cost Effectiveness Ratio (ICER). Each intervention is compared to the next most effective alternative. The strategy is considered dominated when it generates higher costs and lower effectiveness compared to the alterative strategy. Cost effectiveness thresholds are one-time GDP per capita, US$9,660 (≈RM40,000) and three times GDP per capita, RM120,000.

### 2.7 Ethics approval

Ethics approvals were obtained from Pusat Perubatan Universiti Kebangsaan Malaysia (JEP-2016-360) and the Medical Research and Ethics Committee (MREC), Ministry of Health Malaysia (NMRR-16-1341-30856). This study was registered at ClinicalTrials.gov (NC T02862717).

## 3.0 Results

### 3.1 Life years and quality adjusted life years

Table 3 shows the number of calculated LY and QALY. The average LY was 4.15 and 3.70 years for HD and CAPD respectively. Based on EQ-5D-3L index utility scores, average QALY for HD was 3.544 and 3.348 for CAPD.

**Table 3:** Cost effectiveness and cost utility analysis

| Costs and outcomes | HD | CAPD |
| --- | --- | --- |
| Life year (LY) | 4.15 | 3.70 |
| Quality adjusted life year (QALY)<sup>a</sup> | 3.544 | 3.348 |
| Cost per Life year (RM)<sup>b</sup> | 39,791 | 37,576 |
| Cost per QALY (RM) | 46,595 | 41,527 |

<sup>a</sup>=Mean utility index for HD (0.854) and CAPD (0.905) [15]
<sup>b</sup>=Mean cost per patient per year, RM39,791 for HD and RM37,576 for CAPD [14]

### 3.2 Cost effectiveness and cost utility of HD and CAPD

The cost per LY for patients on HD was RM39,791, slightly higher than the cost per LY for patient on CAPD (RM37,576). The cost per QALY for patient in HD was RM46,595 and RM41,527 for patient in CAPD. The cost ratio of HD to CAPD per LY and per QALY was 1.06 and 1.12 respectively (Table 3).

### 3.3 Transitional probabilities

The annual death rate was higher in CAPD (0.134) than in HD (0.125). CAPD patients had a higher rate of switching dialysis modality (0.067) than HD patients (0.007) (Table 4).

**Table 4:**
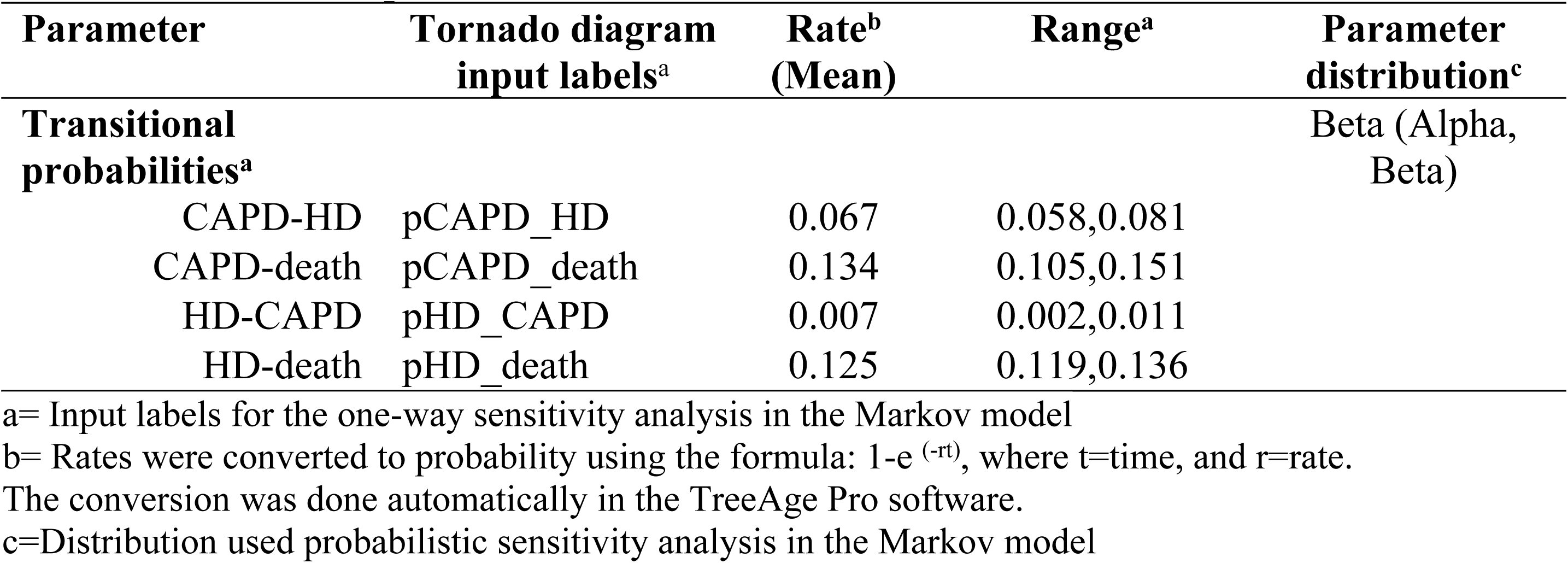
Transitional probabilities

### 3.2 Markov model

#### 3.2.1 Projected costs, outcomes and cost effectiveness

Table 5 shows the results of the Markov model cohort simulation. Scenario 1 (55% HD and 45% CAPD) and scenario 3 (70% HD and 30% CAPD) were dominated strategies. The total undiscounted projected costs in scenario 2 were RM307,014 with 7.902 LYs and 7.041 QALYs. The base case scenario generated a higher undiscounted LYs (8.005) and QALYs (7.113) but with a higher cost (RM313,412). The ICER did not exceeded cost effectiveness threshold of three times GDP (RM120,000). However, the ICER exceeded the threshold for discounted costs and outcomes. Thus, scenario 2 appeared to be the most cost-effective strategy.

**Table 5:**
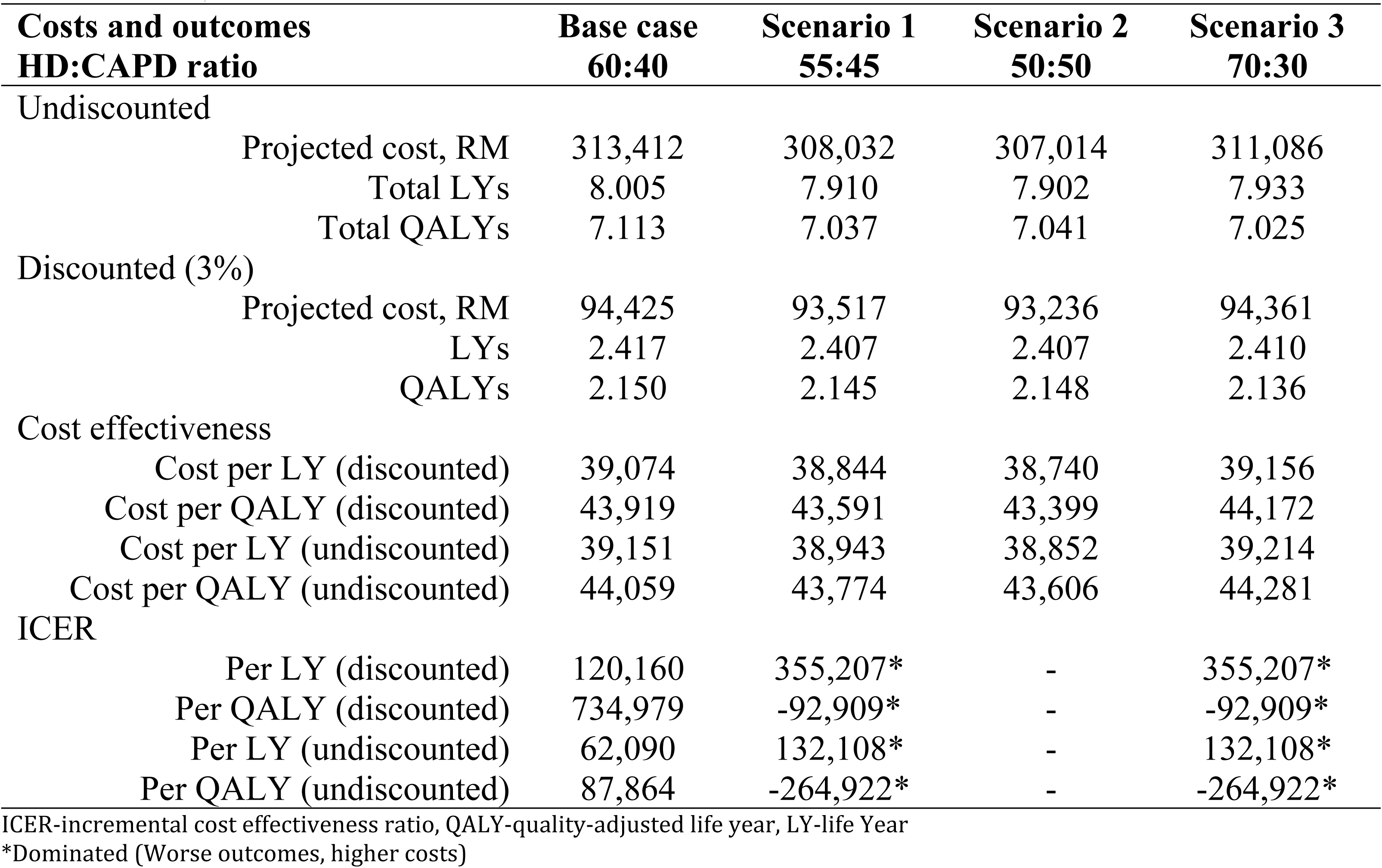
Costs, outcome and cost effectiveness

#### 3.2.2. One-way sensitivity analysis

Figure 2 and Figure 3 show the Tornado diagram with discounted costs and outcomes and undiscounted costs and outcomes respectively. In both sets of results, all imputed values are cost-effective at the cost effectiveness threshold (RM120,000). Health utilities, costs of hospitalizations and costs of outpatient clinic care in both modalities were the top predictors for the uncertainty of effectiveness in the Markov model.

**Figure 2:**
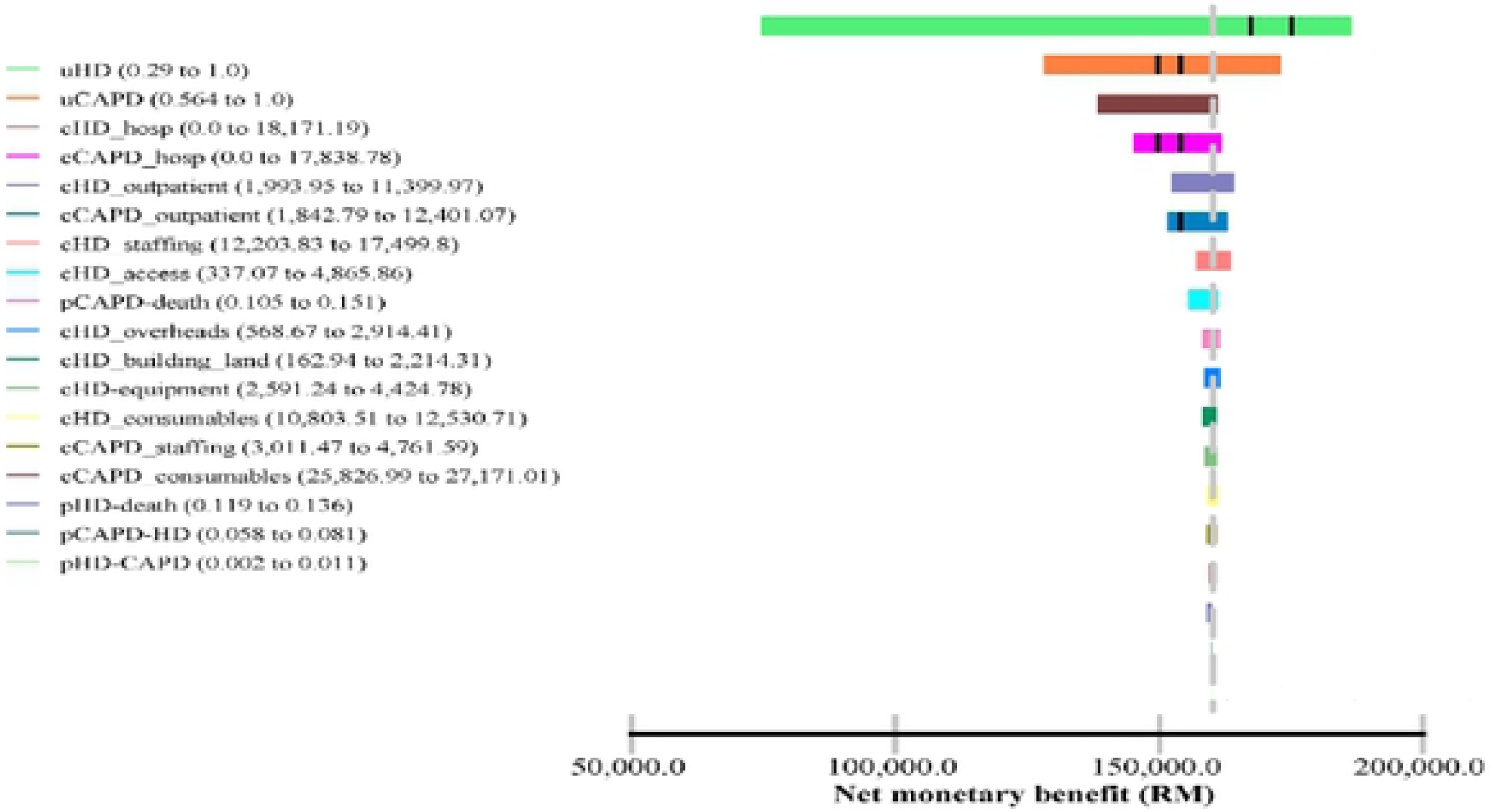
Tornado diagram (discounted) *Cost effectiveness threshold=RM120,000

**Figure 3:**
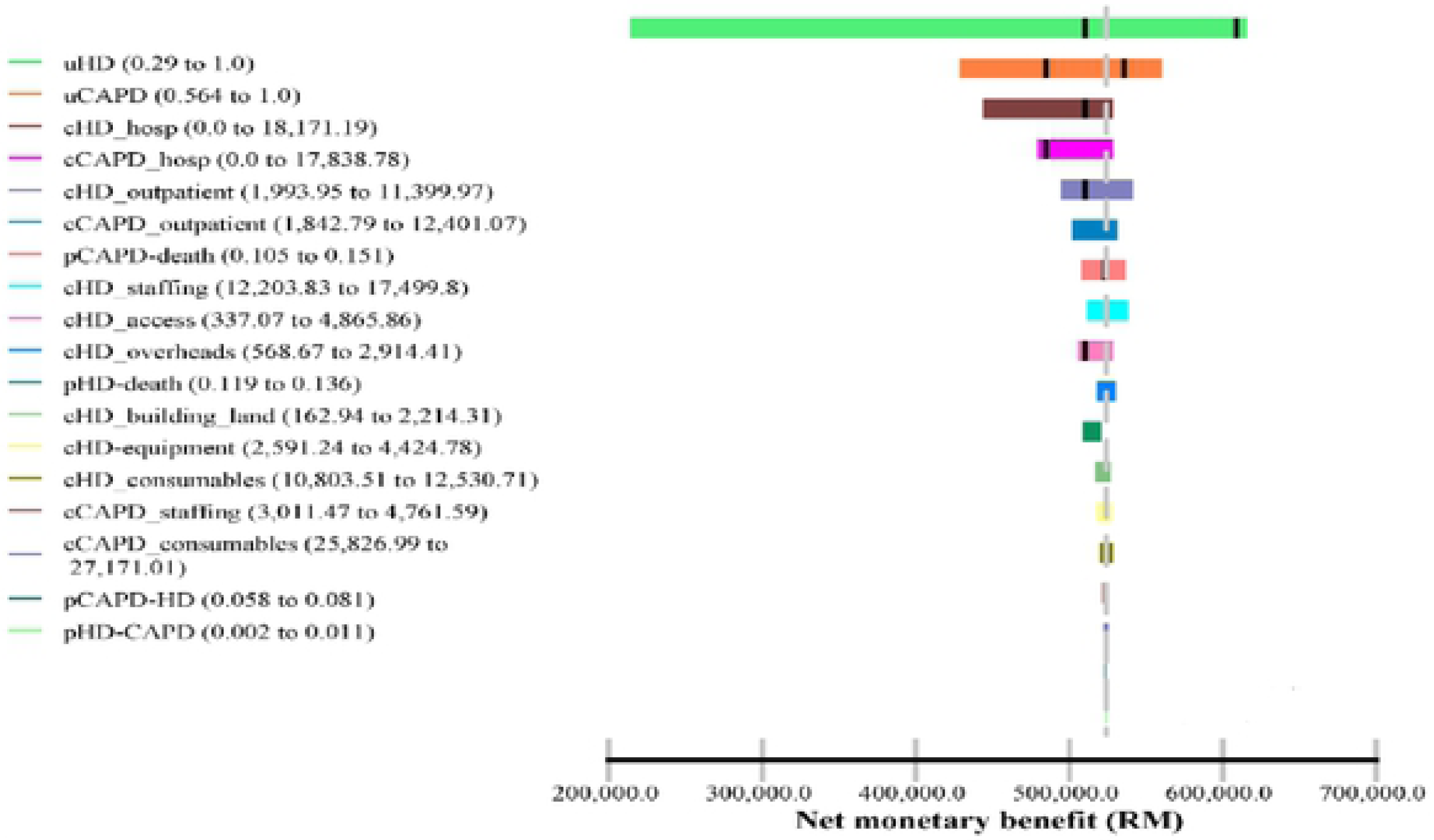
Tornado diagram (undiscounted) *Cost effectiveness threshold=RM120,000

#### 3.2.3 Probabilistic sensitivity analysis

The CEAC of the Markov model (Figure 4) indicates that the probability of favouring base case or Scenario 2 is dependent on the level of the cost effectiveness threshold. At GDP of RM40,000-RM90,000, Scenario 2 was the best option. The base case was the best option if the accepted threshold is more than RM90,000. Irrespective of GDP threshold values, Scenario 1 and Scenario 3 were not cost-effective.

**Figure 4:**
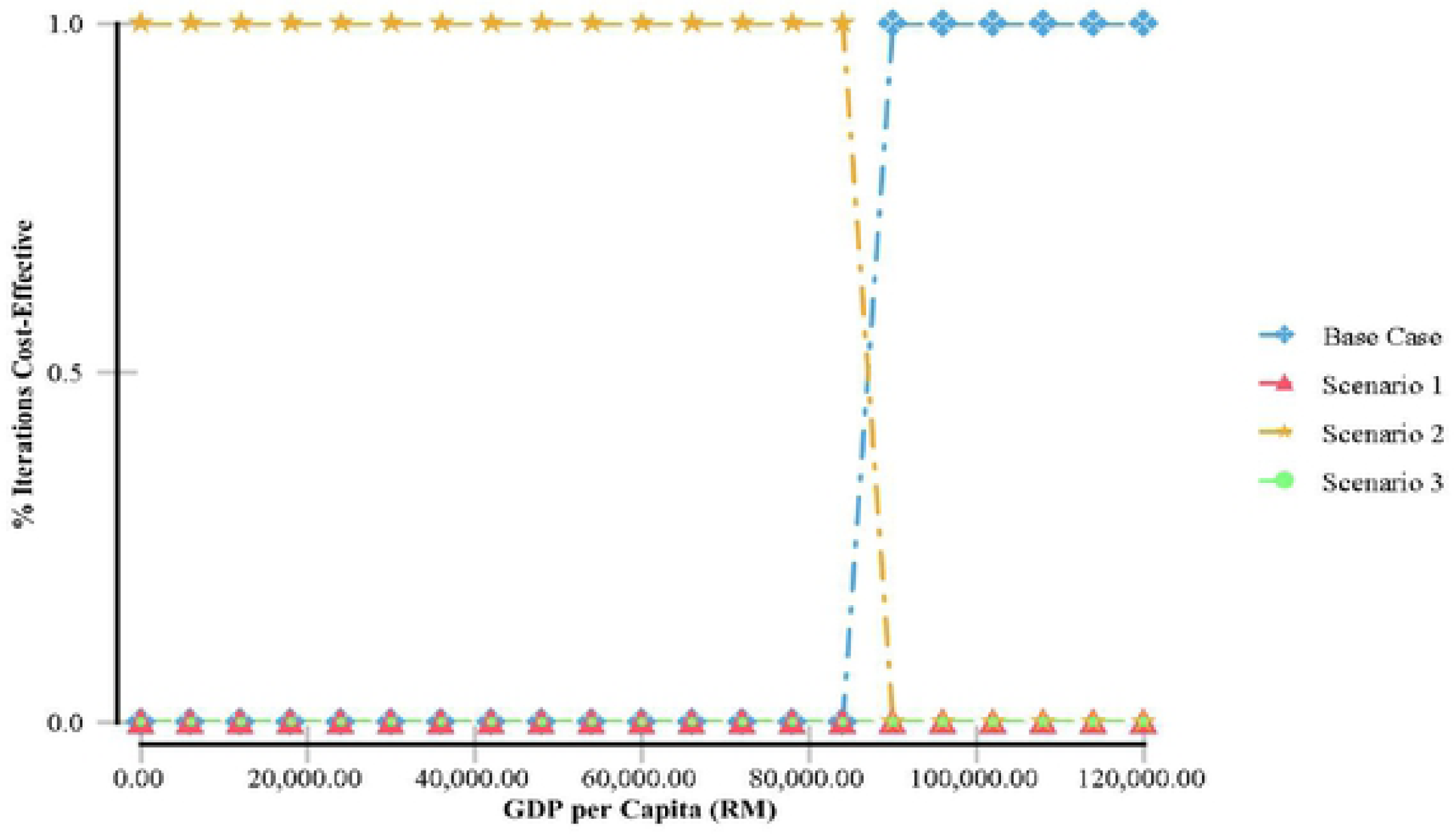
Cost effectiveness acceptability cure (discounted and undiscounted)

### 4.0 Discussion

This cost utility analysis study has provided a cost-analysis framework (micro-costing and step-down approach) and robust results of cost effectiveness of HD and CAPD in Malaysia. This is the first cost utility analysis of dialysis treatments for ESRD patients in Malaysia. The results indicate that CAPD is slightly more cost-effective than HD and the results are consistent with the previous economic evaluation of HD and CAPD in MOH centres in Malaysia [20].

However, the difference of costs per QALY or LY between HD and CAPD was small and not comparable to most developed and some developing countries [2, 21-24]. The ratio of HD to PD costs ranged from 0.70 in Nigeria to 1.90 in Canada [21]. The comparison of costs between HD and PD is presented in ratio forms to avoid possible biases introduced by heterogeneity in currency, eliminating the need for conversion rates and adjusting for inflation rate [21]. They highlighted that HD is generally more expensive than PD in developed countries, but data was not adequate to make any generalizations about the costs in developing countries. In developed countries, due to expensive labor and infrastructure costs, HD is frequently reported to be more expensive than CAPD [2]. For instance, Singapore has a 1.38 HD to PD cost ratio and the PD fluid is manufactured locally [24]. Just et al. (2008) reasserted their view that in developing countries where there are inexpensive labor costs and high imported equipment and solution costs, PD is more expensive than HD [2]. In Malaysia, the main cost component of HD is labor costs while dialysis consumables contribute a significant portion of total costs for CAPD [14]. The LYs and QALYs were higher in HD than in CAPD. The difference of survival between HD and CAPD may not be directly due to the dialysis modality. Survival rates are confounded by clinical and non-clinical factors [25-30]. In Malaysia, the apparent difference of the mortality risk between HD and CAPD is partly attributed to negative selection of PD patients [11]. The lesser LYs gained on CAPD was not compensated by a large increase in health utilities. Unlike in other countries utilities did not differ significantly in Malaysia [15]. In addition, the cost per QALY for both modalities exceeded RM40,000 which implies that both modalities are not highly cost-effective. This does not reflect the true scenario since Malaysia is a country where the cost per QALY is low and the GDP is increasing yearly. Quoting the International Monetary Fund, GDP per capita for Malaysia rose from US$4,290 in 2000 to US$9,660 in 2017. Another important factor to consider in interpreting the results is that, the value of Ringgit Malaysia dropped significantly in the past few years with the lowest in a decade (US$1=RM4.54) recorded in November 2016. Although the value of RM improved in 2017, it was still very low, average US$1=RM4.30.

The Markov model is an analytical framework that is often used in decision analysis and is possibly the most common type of model used in economic evaluation studies [31]. Markov models are a popular form of decision-analytic model which distinguish patient cohorts based on a finite number of mutually exclusive “health states”. The Markov model in this study shows that Scenario 2, 50% HD and 50% CAPD is the most cost-effective strategy. Scenario 2 incurred lesser costs but marginally lesser effectiveness than the base case scenario (60% HD and 40% CAPD). However, the ICER for the base case exceeded one-time GDP and three times GDP for undiscounted and discounted respectively. The Markov model is the first attempt to examine the cost utility of the different strategies of the dialysis provision in Malaysia.

The findings are consistent with the results reported by several countries on this topic in terms of PD expansion. The Markov model conducted by Treharne et al. (2014) analyzed the incident dialysis population to determine whether the proportion of patients on PD should be increased in United Kingdom. Compared with the reference scenario (22% PD, 78% HD), increasing PD use (39 % PD, 61% HD) and (50% PD, 50% HD) resulted in reduced costs and better outcomes. Both strategies dominated the third scenario (5% PD, 95% HD) [32]. The study by Howard et al. (2009) in Australia reported that starting 50% of patients commencing RRT on PD resulted in significant cost savings and was at least as effective as the base case (12.5%) [33]. Similar observations were reported in Austria [34], Spain [17], Norway [35] and Indonesia [36]. In a budget impact analysis in Malaysia increasing PD provision contributes to cost savings. It will improve patients’ access to dialysis in rural areas of Malaysia as the current funding model favours the setting up of HD centres in urban areas [37].

In the present study, an increased 5% CAPD uptake is still a dominated scenario. In contrast, the Markov model developed by those countries mentioned above, showed favourable effectiveness and cost effectiveness in all scenarios when CAPD proportion is increased. This situation can be explained by several reasons. There is an apparent advantage of the mortality rate for HD in the current Markov model. In the other Markov models, PD had lower death risk than HD (the survival advantage favours PD). In countries where demographic and comorbidity data was comparable in both groups of patients, the disadvantage of survival on PD was not observed. Some countries adopt propensity cross matching approach to compare the relative effectiveness of both modalities. In such attempt by Chang et al. (2016), they postulated that the estimated life expectancy between HD and PD were nearly equal (19.11 versus 19.08 years) in the national cohort study with 14 years follow-up [25]. However, propensity score and adjustments were not pursued in the current study to reflect the current situation in Malaysia. Hence, the unadjusted mortality rate was higher in PD than HD in the current Markov model.

There is low technique survival in PD patients in Malaysia which means there is a high probability of PD patients converting to HD annually. The rate of CAPD to HD transition used in this model was 6.70% (range 5.80% to 8.10%) annually. The 24^th^ MDTR report stated that one-year PD technique survival was 94% and 66% at five years (censored for death and transplant) [11]. Technique survival is crucial for PD programme expansion alongside other factors such as catheter placement and patients’ education [38]. In contrast, HD patients enjoy excellent technique survival in Malaysia. The one-year HD technique survival was 99% and 97% at five years (censored for death and transplant) [11]. Because of the high technique failure in CAPD patients in Malaysia, the HD unit must be prepared to cater for patients who are likely to fail CAPD. Most HD units keep one HD machine free for every 40 CAPD patients on treatment [20]. Another important factor to consider when interpreting the results is the insignificant difference in the cost between HD and CAPD in the current study. Other Markov models heavily favour PD expansion due to the large difference in the costs of dialysis accompanied by the positive effectiveness in PD.

The one-way sensitivity analysis via the Tornado diagram shows that health utilities, hospitalization costs and costs associated with outpatient clinic care relatively have a large impact on the net monetary benefits (NHB). Costs related to staffing, overheads, dialysis consumables, land and building have little to no sensitivity to the NHB. These findings accentuated the uncertainties in the Markov model and probably, the cost effectiveness relies on individual patient’s characteristics. The probabilistic sensitivity analysis via the CEAC, indicates that Strategy 2 (50% CAPD) is very cost-effective strategy. The base case is favourable if the cost effectiveness threshold is accepted in the region of above RM90,000. This would be unlikely considering the mean willingness to pay (WTP) among Malaysian population in one of the states in Malaysia was RM 29,080 (US$9,000) in 2010, per additional QALY gained [39].

The present study has several limitations. The lack of randomized controlled clinical trials means the causality between dialysis modality and mortality cannot be determined. Training costs of dialysis staff was not taken into the consideration in the cost analysis. It is recommended to include training costs in the cost analysis [16]. Kidney transplant was not included as one of the health states in the Markov model. Kidney transplant rate from deceased donors in Malaysia is very low and the annual probability of dialysis patients receiving kidney transplants from deceased donors is minute. The model was also kept simple without sub-group analysis and only the observed rates were used to minimise the complexity of the analysis while ensuring the research objectives were met.

## 5.0 Conclusion

In conclusion, both HD and CAPD are viable dialysis modalities in Malaysia. The Markov model favours CAPD expansion but with limitations. Hemodialysis and CAPD are established dialysis modalities that complement each other. A very important advantage of expanding home-based treatment like CAPD is that patients’ disparities in access to dialysis can be improved particularly in less developed areas. The MOH through numerous agencies is already taking steps to encourage ESRD patients without contraindications to consider CAPD as a treatment option. Although reimbursements, economic considerations and government policies are imperative in dialysis provision, patient’s preference cannot be overlooked. Patient selection is also key to a successful CAPD programme because patient’s technique survival is still a major issue in CAPD.

## Acknowledgements

The authors would like to gratefully acknowledge all the people that have made this study possible. First and foremost, we would like to thank the sub principal investigators and research assistants comprise of nurses and medical assistants at each centres for their valuable input and data collection; Dr Liu Wen Jiun, Ms. Jamilah Sarif, Mr. Norisham bin Mohd Dom (Hospital Sultanah Aminah), Dr Kiren Kaur A/P Bhajan Singh, Ms. Rozana Bt Zainol Rasid, Ms. Bistari Binti Zubir (Hospital Tengku Ampuan Afzan), Mr. Amirul Nizam bin Mohtar, Mr. Mohd Patrizal bin Zahari, Ms. Jamaiyah binti Supar, Ms. Vijaya A/P Lakayan (Hospital Kuala Lumpur), Ms. Lim Siew Kim, Mr. Khairul Nul Hakim bin Hazman, Norhazliza binti Hashim Hospital Pulau Pinang, Mr. Ratneswaran A/L Naganathan, Ms. Noriah binti Othman (Hospital Tengku Ampuan Rahimah). Second, we would like to acknowledge Dato’ Dr. Tan Chwee Choon, former Head of Nephrology Service, Ministry of Health Malaysia and Datuk Dr. Ghazali Ahmad, Head of Nephrology Department, Hospital Kuala Lumpur for providing us with their able assistance and allowing us to conduct this research. Besides, we would like to thank National Renal Registry, in particular, Madam Lee Day Guat in providing patients’ list for sampling. Finally, the authors thank the Director General of Health in Malaysia for permission to publish this paper.

